# Arousal-informed neuronal avalanches operate near criticality

**DOI:** 10.64898/2026.10.12.757995

**Authors:** Cédric Stefens, Hardik Rajpal, Joe S. Canzano, Meghdad Saeedian, Michael G. Kareithi, Henrik Jeldtoft Jensen, Mauricio Barahona, Spencer LaVere Smith, Simon R. Schultz

## Abstract

Cortical networks are thought to operate near a “critical point” that optimizes information processing, but this has been difficult to test at single-cell resolution across large brain areas, as narrow fields of view miss overlapping neural avalanches, causing subcritical estimates, Using mesoscopic two-photon calcium imaging in awake mouse visual cortex, we recorded thousands of neurons and applied convolutional non-negative matrix factorization to decompose population activity into distinct, overlapping spatiotemporal avalanches. Rather than occupying a single dynamical state, we find that the cortex hosts multiple overlapping subnetworks distributed across a phase diagram at varying distances from the critical point. Arousal-linked avalanches propagate widely and operate nearest the critical point, showing optimized dynamic range, minimized algorithmic complexity, and maximized temporal entropy. Our results reveal how the awake brain tunes parallel near-critical subnetworks to support robust sensory computation.

## Introduction

The brain continuously integrates external sensory information with internal cognitive states to generate adaptive behaviour. To maintain both stability and flexibility, theoretical and empirical evidence suggests that cortical networks self-organize near a critical phase transition between ordered (synchronized) and disordered (asynchronous) states (Beggs & Plenz, 2003; Chialvo, 2018). Operating at or near this critical point is hypothesized to optimize computational properties, including information capacity, the fidelity of information transmission, and dynamic range (Shew et al., 2009; Shew & Plenz, 2013). The hallmark of this dynamical regime is the emergence of scale-free, spontaneous cascades of neural activity known as neuronal avalanches. In idealized in vitro and anesthetized in vivo preparations, these avalanches characteristically exhibit power-law distributions with a size exponent of τ ≈ 1.5 and a duration exponent α ≈ 2.0, aligning with the universality class of a mean-field directed percolation or critical branching process (Beggs & Plenz, 2003; Friedman et al., 2012; Plenz et al., 2021).

Despite extensive theoretical support, characterising criticality in the awake, behaving brain has proven challenging. Measurements of cortical dynamics in awake animals frequently reveal deviations from ideal power-law scaling, often pointing toward weakly subcritical regimes (Priesemann, 2014; Wilting & Priesemann, 2019). These discrepancies stem largely from inherent methodological limitations in recording and analysis. Conventional avalanche detection algorithms assume that cascades occur as isolated, sequential events separated by quiescent periods. However, large-scale population recordings indicate that awake cortical dynamics are high-dimensional and simultaneously driven by multiple sequences of activity (MacDowell & Buschman, 2020; Stringer et al., 2019). When overlapping avalanches are forcibly segregated by rigid temporal binning, or when networks are severely spatially subsampled due to restricted fields of view (FOV), the resulting distributions systematically misrepresent the underlying cascade dynamics and flatten the tails of the probability distributions (Priesemann, 2014; Srinivasan et al., 2024). Consequently, a gap remains in our understanding of how scale-free dynamics can be measured in high-dimensional large-scale recording methods where temporal and spatial subsampling persist.

Recent analyses suggest that critical dynamics may not govern the entire macroscopic network, but instead be embedded within specific low-dimensional subspaces of cortical activity (Fontenele et al., 2024). Furthermore, cortical state is not static; it undergoes rapid fluctuations driven by neuromodulation and arousal, which dynamically alter the balance of excitation and inhibition (Reimer et al., 2014; Waschke et al., 2019). Brain-wide propagating waves and spontaneous neural cascades are closely coupled to autonomic indices of arousal, such as pupil diameter, optimizing sensory encoding and behavioural responsiveness (Raut et al., 2021, 2025; Yang et al., 2026). This raises the possibility that the cerebral cortex does not reside at a single fixed distance from criticality, but rather supports multiple concurrent, overlapping subnetworks that are differentially tuned toward or away from the critical point by fluctuations in arousal and sensory drive.

Arousal here refers to the level of neural and physiological activation influencing responsiveness, vigilance, and sensory processing, rather than being a synonym for locomotion or stimulus presentation (McCormick et al., 2020; Joshi and Gold, 2020). Pupil diameter is a useful but non-specific arousal index: its fluctuations also reflect luminance and other visual and behavioural influences (Ruuskanen and Mathôt, 2026).

We tested the hypothesis that overlapping cortical subnetworks differ in their proximity to criticality according to their coupling to arousal. We used mesoscopic two-photon calcium imaging to record from thousands of neurons across a wide FOV in the awake mouse visual cortex, encompassing both striate and extrastriate areas. To overcome the limitations of conventional, single-event avalanche detection, we applied convolutional non-negative matrix factorization (seqNMF) (Mackevicius et al., 2019) to decompose continuous population activity into distinct, co-occurring spatiotemporal sequences. Avalanches detected within each reconstructed sequence, termed sequence-avalanches, were mapped onto an order–control phase diagram relating population activity (order parameter) to the estimated branching parameter (control parameter). We then examined whether sequence-avalanches with a stronger coupling to pupil-related arousal were associated with greater proximity to criticality and enhanced information-processing metrics.

## Results

### 1. Neuronal avalanches in the striate and extrastriate visual cortical areas layer 2/3 operate near the critical point

The theory of self-organized complex systems suggests that cortical circuits operate near a critical phase transition, an optimal regime for information processing (Beggs, 2008; Beggs & Plenz, 2003; Shew et al., 2009, 2011; Shew & Plenz, 2013) To investigate this framework at the mesoscopic scale (across millimetres of cortex), we hypothesized that the sensory-stimulated state of the brain should be actively tuned to be closer to the critical point in comparison to the spontaneous state, thereby maximizing the network’s dynamic range and the repertoire of accessible states available to process incoming stimuli.

Mesoscopic calcium imaging of layer 2/3 populations across a 3x3 mm² field of view (FOV) in the awake mouse brain enabled the identification and characterization of widespread neuronal avalanches (Fig. 1d). We recorded from head-fixed mice (n=8 animals; 10 recordings) across two primary environments: a spontaneous condition (dark or grey screen, n=5 recordings) and a sensory-driven condition (randomized gratings or naturalistic scenes, n=5 recordings). Because the high cell count (4736 ± 1052 neurons for spontaneous and 7242 ± 606 neurons for sensory datasets) yields a non-zero and uninterrupted summed activity (Fig. 1b), we applied an optimizing-threshold method that segment the activity into discrete events and maximizes the avalanche rate (Fig. 1c) (Bellay et al., 2015; Curic et al., 2021). The optimal binarization threshold (θ) did not significantly differ between the spontaneous (16.11 Hz ± 0.38 SD) and sensory conditions (15.51 Hz ± 0.46 SD; p = 0.331, permutation test).

**Figure 1.**
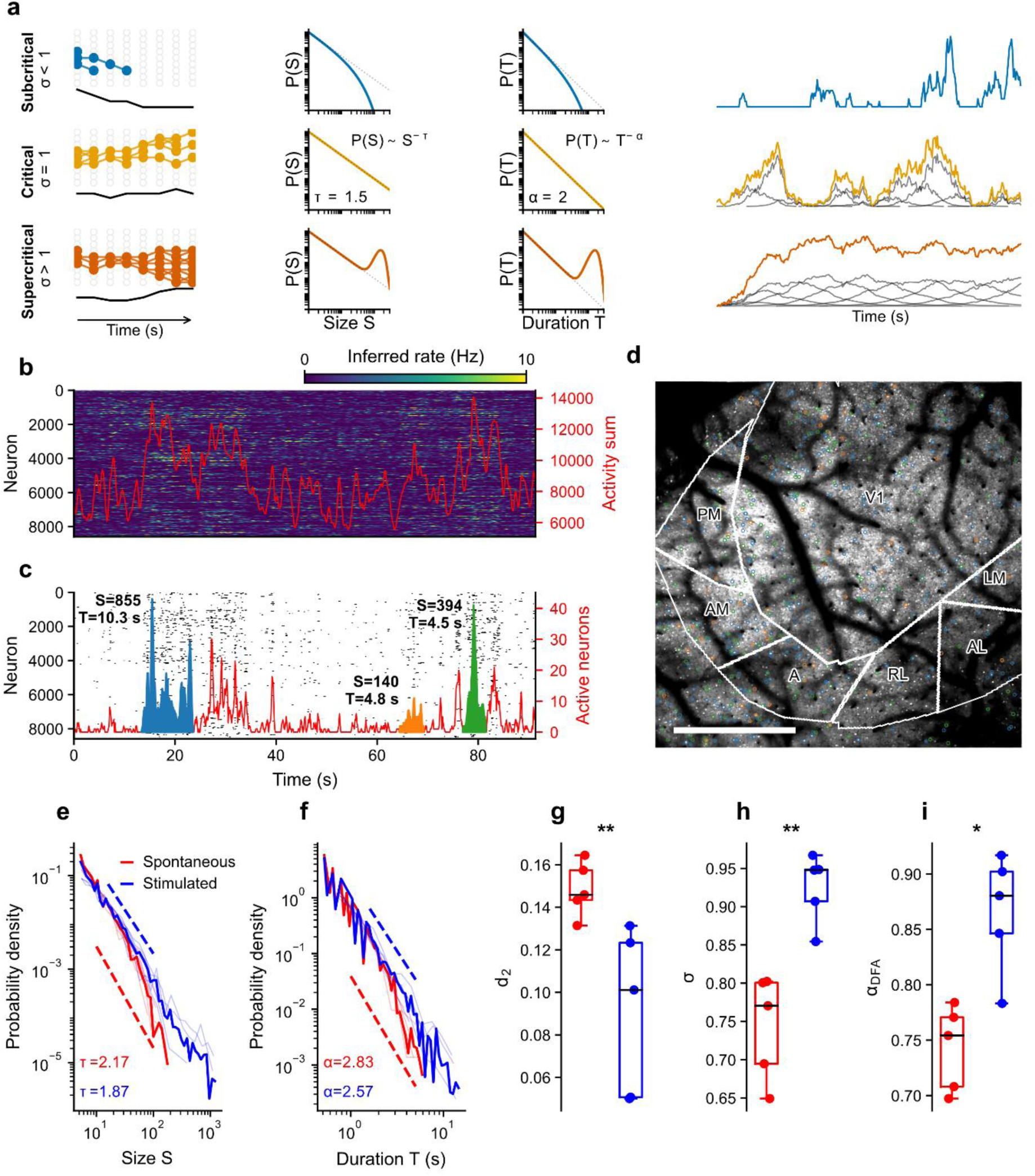
Context-dependent signatures of criticality in the visual cortex layer 2/3. **(a)** Schematic representation of subcritical (branching parameter *σ* < 1), critical (*σ* = 1), and supercritical (*σ* > 1) regimes. Left: Network propagation, black lines represent sum of active cells; Middle: Avalanche size and duration distribution; Right: sum of active neurons time series, underlying overlapping of single avalanches depicted in the left panel. **(b)** Raster plot of deconvolved calcium activity (spiking rate) for a single animal (90 s excerpt). The red curve denotes the population sum activity. Continuous activity prevents the separation of clusters into distinct avalanches without thresholding. **(c)** Binarized raster plot (black) using an optimal threshold *θ* to maximize avalanche detection. Avalanches (colored areas) are defined as excursions above the threshold, characterized by size *S* (number of spikes) and duration *T* (number of time bins). Three example avalanches are highlighted in blue, orange, and green. **(d)** Average mesoscopic image of the primary (V1) and secondary visual cortices. Colored circles indicate neurons participating in the three example avalanches shown in (c). Abbreviations: PM: posteromedial; AM: anteromedial; A: anterior; RL: rostrolateral; AL: anterolateral; LM: lateromedial. **(e)** Avalanche size distributions *P*(*S*) for spontaneous (red) and sensory (blue) conditions. **(f)** Avalanche duration distributions *P*(*T*). Thick lines indicate the average across recordings (*n* = 5 per condition); thin lines represent individual datasets. Dashed lines indicate power-law fits. **(g)** Proximity to criticality (*d*_2_), defined as the distance between the fitted VAR model of the population activity from the hyperplane of temporally renormalizable solutions (lower values indicate closer proximity to criticality) **(h)** Estimated branching parameter (*σ*). A value of 1 indicates criticality. **(i)** Detrended Fluctuation Analysis (DFA) exponent (*α*). A value of 0.5 indicates uncorrelated noise, while ∼ 1.0 indicates long-range temporal correlations (self-similarity). Boxplots show median and interquartile range; points represent individual recordings. ∗ *p* < 0.05, ∗∗ *p* < 0.01 (permutation test).

Using this approach, we detected 23,999 neuronal avalanches. Their frequency remained stable across behavioural contexts (2603.6 ± 158.3 avalanches per session in spontaneous vs. 2196.2 ± 435.5 in sensory; p > 0.05). However, the spatial scale of the network events was highly context-dependent. Avalanches in the sensory condition recruited significantly larger coordinated ensembles (mean size ⟨S⟩ = 15 ± 4 neurons) compared to those in the spontaneous condition (⟨S⟩ = 6.5 ± 0.3 neurons; p=0.040). The average duration (⟨T⟩) did not differ significantly between spontaneous (0.48 ± 0.03 s) and sensory (0.62 ± 0.06 s) states (p=0.078).

Analysis of the probability distributions for avalanche size (S) and duration (T) revealed scale-approximate invariant power-laws with exponential decay at the tail in both conditions, a hallmark of critical dynamics (Fig. 1a, e, f). The sensory condition yielded critical exponents closer to the theoretical mean-field values of τ ≈ 1.5 and α ≈ 2.0 (Beggs & Plenz, 2003). Specifically, the sensory state exhibited a size exponent of τ = 1.87 (Fig. 1e) and a duration exponent of α = 2.57 (Fig. 1f), whereas the spontaneous state displayed steeper decays with τ = 2.17 and α = 2.83.

To rigorously quantify the network’s dynamical regime and to read signatures of near-critical states, we computed three independent metrics. First, proximity to criticality (d2), the deviation from temporal renormalizability of the population activity, was significantly higher in the spontaneous condition (0.149 ± 0.006) compared to the sensory condition (0.091 ± 0.017; p=0.008) (Fig. 1g) (Sooter et al., 2025). Second, the branching parameter (σ), calculated using a multistep regression estimator to account for subsampling (Spitzner et al., 2021), approached the critical value of 1 during sensory stimulation (σ = 0.925 ± 0.020) but indicated a more subcritical regime during spontaneous activity (σ = 0.743 ± 0.031; p=0.003) (Fig. 1h). Finally, Detrended Fluctuation Analysis (DFA) revealed stronger long-range temporal correlations in the sensory state – a hallmark of criticality; the DFA exponent (α_DFA) was significantly higher during sensory processing (0.866 ± 0.024) compared to the spontaneous state (0.743 ± 0.017; p=0.015) (Fig. 1i).

In summary, while the visual cortex maintains a spontaneous state with signatures of criticality, sensory engagement actively tunes the global network dynamics closer to the critical point. However, this full-FOV conventional thresholding approach restricted to high-activity event shows few active neurons in the avalanches, which obscures the underlying heterogeneity of the network activity. To determine whether these macroscopic signatures arise from a uniform critical state or a confluence of functionally specialized subnetworks, it is necessary to decompose the continuous spatiotemporal activity into independent, overlapping components.

### 2. Decomposed neuronal avalanches behave at different distance to criticality along a gradient of pupil mutual information

A recent study suggests that scale-free, critical dynamics may not necessarily be a uniform, global network property, but rather may be confined to specific low-dimensional subspaces co-existing within high-dimensional population activity (Fontenele et al., 2024). To unveil hidden avalanches that utilize sub-threshold neuronal activity discarded by conventional detection methods, we hypothesized that multiple, temporally overlapping neuronal avalanches can occur simultaneously, sharing neurons to propagate. We further posited that the specific characteristics of these decomposed avalanches would predict their distinct proximities to the critical point.

To isolate these underlying subnetworks, we applied convolutional non-negative matrix factorization (seqNMF) (Mackevicius et al., 2019) to the previously analysed datasets (n=10). This dimensionality reduction algorithm factorized the neural activity matrix into distinct spatiotemporal sequences (of ≤ 1.3 seconds) and their corresponding temporal activation amplitudes (Fig. 2.a). The algorithm identified an average of 6-7 distinct spatiotemporal sequences per dataset, which captured the majority of the structured temporal dynamics, accounting for 0.69 ± 0.12 of the total variance in the original data. Convolving these matrices reconstructed overlapping datasets of activity from which 62 valid sequence-avalanches were detected using the same method as in section 1 (Fig. 2.b). Compared to the conventional full-FOV method, the seqNMF avalanches approach explains more data with the number of active cells per time bin raised hundred-fold, reaching into the thousands (Fig. 2.b).

**Figure 2.**
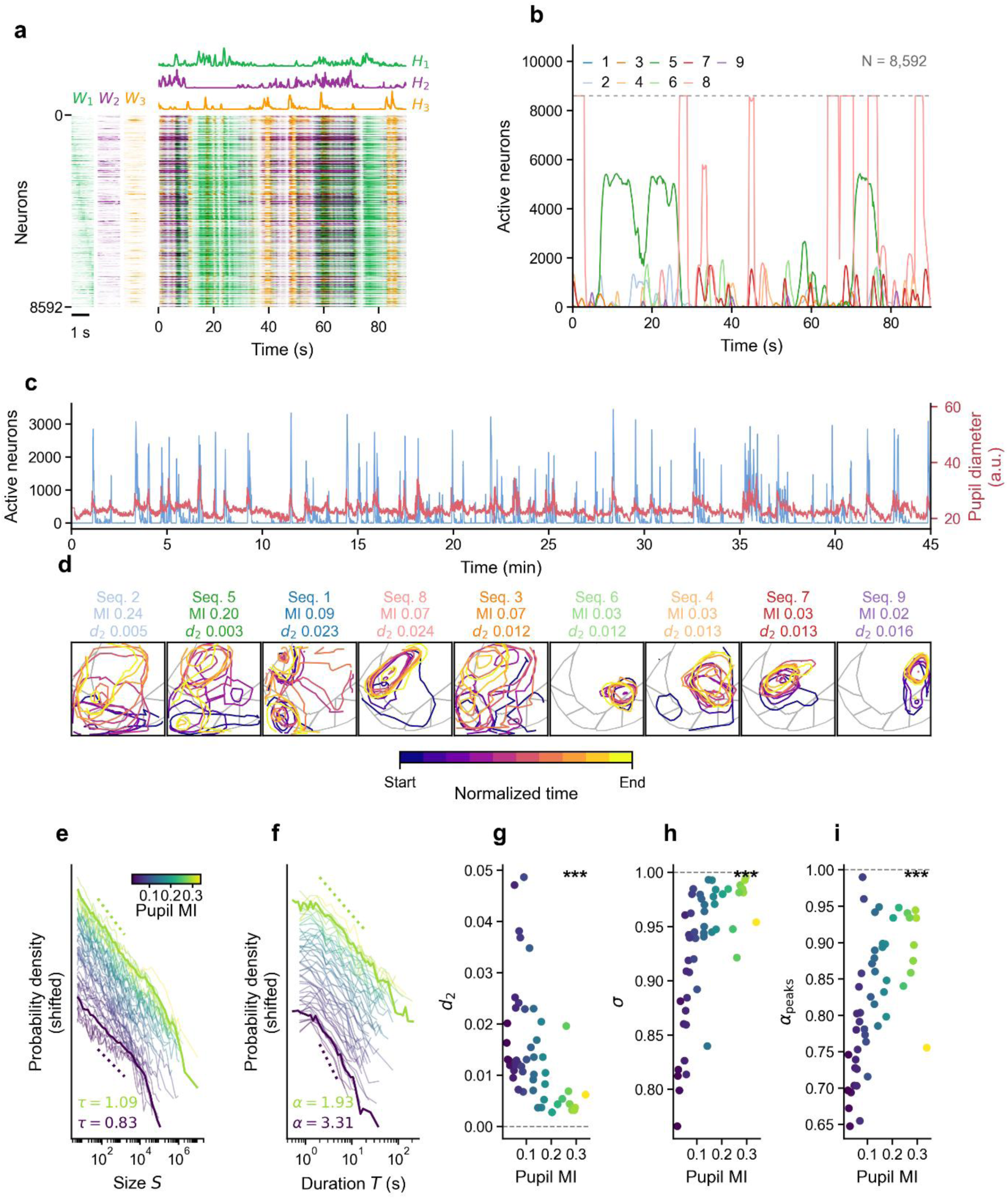
Decomposition into sequences reveals that arousal-informed avalanches operate close to criticality. **(a)** Unsupervised decomposition of mesoscopic calcium imaging data using seqNMF into spatiotemporal motifs (W) and their corresponding temporal weightings (H). **(b)** Activity of nine identified sequence-avalanches (seq. 1–9) on the same segment as in figure 1 b and c. The dashed line marks the population ceiling. **(c)** 45-minute recording of a pupil-informed sequence-avalanche (seq. 2, blue) and pupil diameter (red), demonstrating high temporal coordination. **(d)** Spatiotemporal footprints of the nine sequence-avalanches, ordered by their mutual information (MI) with pupil diameter. Colors indicate normalized time from start (purple) to end (yellow) of the sequence motif. **(e, f)** Probability density distributions for avalanche size (S, panel e) and duration (T, panel f) for individual sequences, color-coded by their Pupil MI. Solid lines indicate the best power-law fits, with exponents (τ and α) computed on sequence-avalanches below the 15^th^ percentile and above the 85^th^ percentile of MI pupil values. **(g-i)** Proximity to criticality as a function of arousal. Scatter plots show (g) proximity to criticality (d2), (h) branching parameter (σ), and (i) DFA exponent (α peaks) plotted against Pupil MI for each identified sequence. Higher pupil-informed sequences demonstrate significantly closer proximity to critical dynamics (lower d2, σ≈1, and α≈1). Statistical significance (***) denotes p<0.001 as determined by a Spearman’s rank correlation test.

To investigate the correlates with arousal of these spatiotemporal sequences, we computed the mutual information (MI) between the sequence-avalanche timeseries and pupil diameter, a well-established proxy for autonomic arousal (Raut et al., 2021, 2025; Reimer et al., 2014, 2016; Ruuskanen & Mathôt, 2026) (Fig. 2.c). Across the 50 sequences with concurrent pupil recordings, MI scores exhibited a broad distribution (mean = 0.13 ± 0.09; range: 0.02 to 0.34). This wide variance indicates a heterogeneous coupling to arousal: while some sequences covaried strongly with pupil dynamics, a substantial fraction operated independently. Ordering the sequence spatial patterns by MI revealed distinct topographical behaviours (Fig. 2.d); high MI avalanches exhibited broad, propagating patterns (some traveling from PPC/S1 to V1) whereas some low MI avalanches remained highly localized within V1, potentially indicating retinotopically organized stimulus responses.

The decomposed avalanche probability distributions showed that arousal-informed avalanches express critical exponents closer to the theoretical values of a mean-field branching process than less informed avalanches. Size distributions plotted as power laws (Fig. 2.e) showed that the average of the 15% highest MI avalanches yielded a size exponent of τ=1.1, approaching the expected 1.5, whereas the average below the 15th percentile exhibited a shallower slope of τ=0.84. Similarly, duration distributions (Fig. 2.f) for high MI avalanches displayed a duration exponent of α=1.93, closely mirroring the expected value of 2.0, while low MI avalanches considerably deviated (α=3.31). Notably, this decomposition approach successfully captured sizes reaching 10^7^ neurons and durations up to 100 seconds, substantially expanding the scale of observation compared to the full FOV method.

Beyond power-law scaling, three independent dynamical metrics confirmed that arousal-informed avalanches operate closer to criticality. First, the proximity to criticality metric (d2) (Sooter et al., 2025) demonstrated that avalanches with higher pupil MI were significantly closer to the temporally renormalizable solutions (Spearman, p<0.001; Fig. 2.g). Second, the branching parameter (σ), estimated via multistep regression (Spitzner et al., 2021), scaled positively with MI, indicating that high arousal avalanches operate near the critical boundary of σ≈1 (Spearman, p<0.001; Fig. 2.h). Finally, Detrended Fluctuation Analysis (DFA) applied to point-process converted event timeseries corroborated these findings, showing that the DFA scaling exponent was positively correlated with pupil MI (Spearman, p<0.001; Fig. 2.i).

The association between pupil MI and d₂ was present separately in stimulated and spontaneous recordings (Spearman rₛ = −0.478, p = 0.0087, n = 29 sequences; and rₛ = −0.696, p = 0.00046, n = 21, respectively). Stimulus MI was available for 29 retained sequences from four stimulated recordings. Stimulus MI showed a weak negative association with event-peak DFA (rₛ = −0.383, p = 0.040), but no significant association with d₂ (rₛ = −0.148, p = 0.444) or branching parameter (rₛ = −0.273, p = 0.152; fig. S2). Individual size and duration fits are shown in fig. S1.

Dimensionality reduction reveals that cortical dynamics are driven by a confluence of temporally overlapping subnetworks operating at varying distances from the critical point. Spatiotemporal avalanches tightly coupled to fluctuations in arousal state operate in a near-critical regime, exhibiting propagating spatial patterns and scale-free exponents that tend to match theoretical predictions. Having established that these specific arousal-informed sequence-avalanches operate near criticality, the subsequent analysis examines whether these near-critical sequence-avalanches optimize neural computation and information processing as predicted by theory.

### 3. Criticality enhances computational capability

The criticality hypothesis predicts that neural networks operating near a critical phase transition can exhibit enhanced dynamic range, information transmission, and information capacity (Shew et al., 2009, 2011; Shew & Plenz, 2013). Having established that decomposed sequence-avalanches can be mapped along a gradient of distance to criticality (d2), we sought to evaluate whether this tuning relates to information processing metrics.

We investigated the relationship between the proximity to criticality (d2) and the computational properties of the 62 demixed sequence-avalanches. Correlation analyses revealed that information storage, which quantifies how much information the past of a sequence-avalanche activity time series provides about its next state, increased significantly as avalanches approached the critical point (Spearman, p < 0.001; Fig. 3a). This suggests that near-critical sequence-avalanche dynamics retain more information about past activity, potentially supporting temporal integration. Furthermore, the dynamic range of the subnetworks was optimized near criticality, demonstrating a strong negative correlation with d2 (Spearman, p < 0.001; Fig. 3.b). This expansion of dynamic range provides evidence that near-critical avalanches are primed to encode more refined and diverse sensory information.

**Figure 3.**
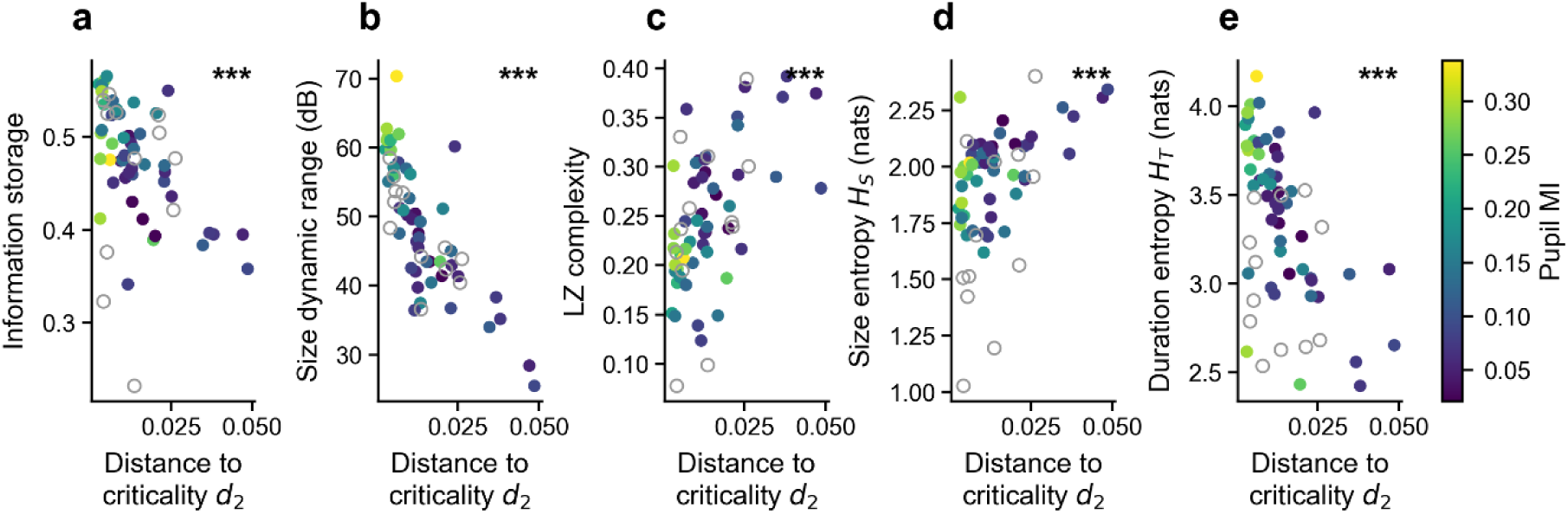
Proximity to criticality optimizes neuronal avalanches computational capacity and information richness. Relationship between proximity to criticality (d2) and information-theoretic metrics for sequence-avalanches. **(a)** Information Storage, quantifying the network’s ability to maintain information over time. **(b)** Dynamic range, the span of distinguishable intensities occurring in each sequence-avalanche. **(c)** Lempel-Ziv (LZ) Complexity, assessing the richness and non-randomness of temporal occurrences. **(d)** Information entropy of avalanche durations indicating the richness of temporal information. **(e)** Information entropy of avalanche sizes, representing the capacity to encode the magnitude of neural events. Statistical significance (***) denotes p<0.001 as determined by a Spearman’s rank correlation test. Each point represents one sequence-avalanche. Filled circles are coloured by the mutual information between sequence- avalanche activity and pupil diameter (Pupil MI), Open circles indicate sequence-avalanches without pupil recordings.

We further evaluated algorithmic randomness by computing Lempel-Ziv (LZ) complexity. As sequence-avalanches approached the critical boundary, LZ complexity decreased (Spearman, p < 0.001; Fig. 3.c). Because LZ complexity achieves its theoretical maximum in purely random, statistically independent systems (Mediano et al., 2023), the emergence of structured, long-range spatiotemporal correlations near the critical point naturally reduces the unpredictability of the neural signal. This observation aligns with recent theoretical frameworks demonstrating that LZ complexity drops as neural systems transition from disorganized, supercritical states toward the critical point (Singleton et al., 2023). Consequently, near-critical, arousal-informed avalanches appear to trade pure spatial noise for highly structured, coordinated information processing.

Finally, we assessed the Shannon entropy of both avalanche sizes and durations to characterize spatial and temporal repertoires. The spatial entropy of avalanche sizes decreased significantly near criticality (Spearman, p < 0.001; Fig. 3.d). In the sub-critical regime, fragmented network activity lacks long-range spatial coordination; when subjected to two-photon imaging spatial subsampling, this disconnected activity manifests as a disorganized, noisy distribution that inflates spatial entropy. As arousal tunes the system toward criticality, these disconnected events unify into highly structured, scale-invariant avalanches, naturally reducing spatial noise and the corresponding size entropy. Conversely, the temporal entropy of avalanche durations was maximized near the critical point (Spearman, p < 0.001; Fig. 3.e). Sub-critical dynamics restrict the network’s temporal repertoire because cascades are rapidly quenched, resulting in brief, transient bursts and low temporal entropy. Approaching criticality overcomes this quenching, granting the network access to a vast, scale-free repertoire of sustained timescales that significantly expands the temporal state space and optimizes richness of information.

Collectively, these findings provide empirical support for the hypothesis that information processing metrics are optimized near criticality. The data indicate that high pupil mutual information (MI), arousal-informed avalanches operating near the critical point effectively enhance information processing capabilities. Because the critical point may represent a phase transition between distinct system states, the next question is whether these computation- enhanced avalanches systematically populate a control-order phase diagram near the critical point.

### 4. Confluent avalanches populate the order-control phase diagram at various distances from criticality

On a control-order phase diagram, the coordinates of avalanches define a phase line that can reveal either a first-order (discontinuous) or higher-order (continuous) phase transition. We hypothesized that the decomposed sequence-avalanches would populate this phase line at varying distances from the critical point, with arousal-informed avalanches clustering close to it.

The phase diagram in Figure 4a describes how the system evolves from subcritical to supercritical regimes as an order parameter is tuned by a control parameter, such as excitation–inhibition balance or temperature (Muñoz, 2018). Using the estimated branching parameter as the control variable (Spitzner et al., 2021), we plotted sequence-avalanches on the diagram. Across regimes, the resulting spatial patterns closely resembled those of the Ising model (Fig. 4.a, left side, Fig. 2.d), in which localized domains dominate away from criticality, whereas near the phase transition larger, irregular structures emerge. Consistent with this analogy, subcritical states were characterized by spatially confined islands of activity, while near the critical point activity became spatially extended and scale-invariant.

**Figure 4.**
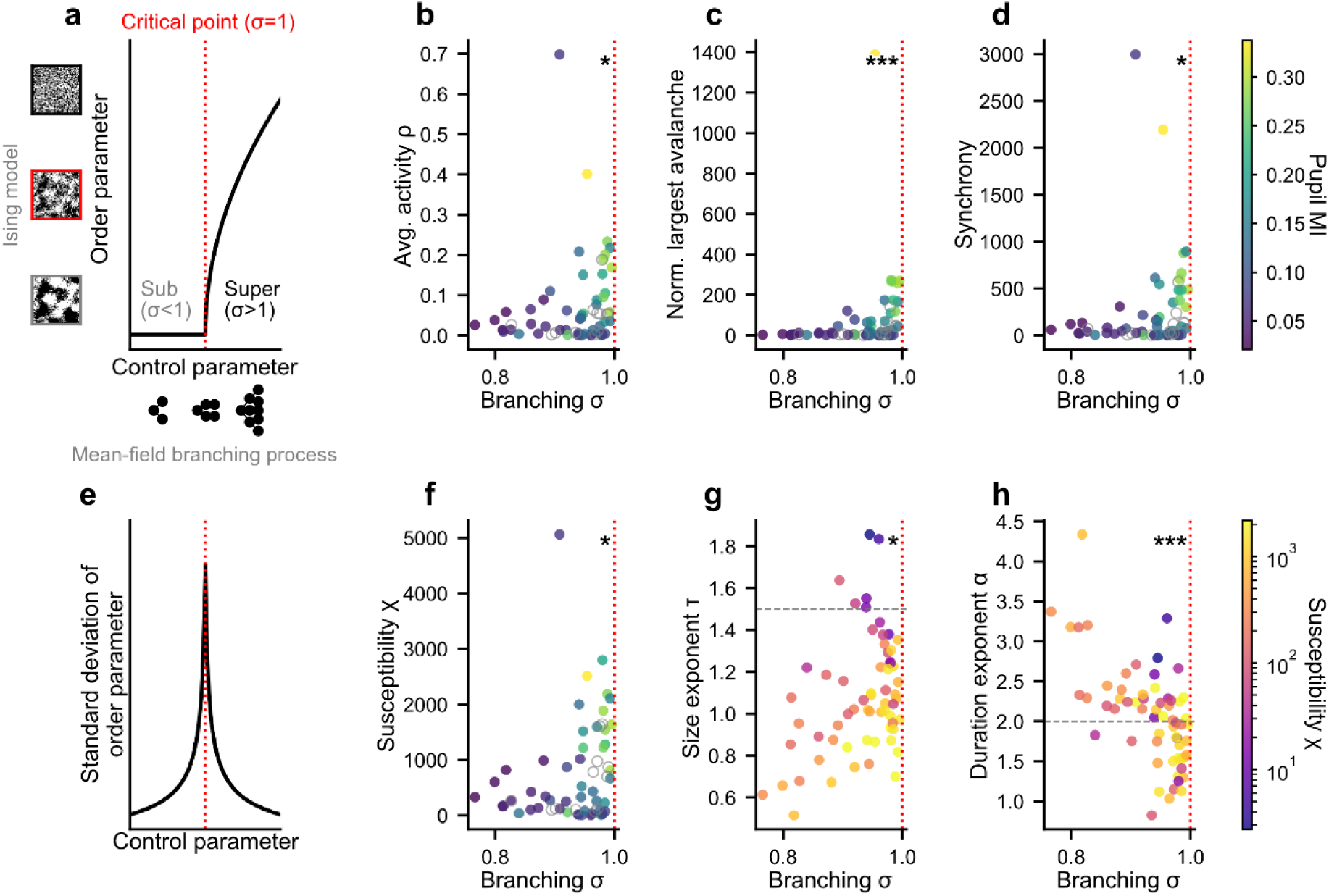
Role-dependent neuronal avalanches populate a universal second-order phase transition line. **(a)** Theoretical schematic of a second-order phase transition (e.g., Ising model or branching process). The critical point (σ=1) separates a disordered subcritical phase from an ordered supercritical phase. Evolution of macroscopic order parameters as a function of the branching parameter (σ): **(b)** Average activity (ρ), **(c)** population-normalized largest avalanche size, and **(d)** network synchrony. These order parameters peak as the system approaches the critical boundary. **(e)** Standard deviation of the order parameter diverges near the critical point. **(f)** Susceptibility (χ) as a function of the branching parameter. The divergence of susceptibility near σ≈1 provides a piece of evidence for a phase transition. Scaling exponent flow: **(g)** The size exponent (τ). **(h)** The duration exponent (α) flows toward the theoretical mean-field value as the network approaches criticality. Statistical significance (∗ *p* < 0.05, ∗∗ *p* < 0.01, ∗∗∗ *p* < 0.001) is determined by Spearman’s rank correlation. Empty circles indicate data without pupil recordings.

To position the avalanches on the diagram, we evaluated the y-axis with multiple order parameters as a function of the branching parameter. Using average activity (ρ) as an order parameter revealed a bifurcation at the critical point, with arousal-informed avalanches situated near this transition (Spearman, p = 0.0351; Fig. 4.b). Similar bifurcations localizing high-MI avalanches to the critical boundary were observed when utilizing the normalized largest avalanche size (Spearman, p < 0.001; Fig. 4.c) and synchrony (Spearman, p = 0.0231; Fig. 4.d) as alternative order parameters.

Theory predicts that susceptibility diverges at a continuous critical point in the infinite-system limit; finite systems instead exhibit a finite peak near the transition (Fig. 4e; Muñoz, 2018). Here, we used a fluctuation-based susceptibility proxy, χ = var(Aₜ)/mean(Aₜ), where Aₜ denotes sequence-avalanche population activity. Consistent with this prediction, susceptibility (χ) exhibited marked increase near the critical point, with arousal-informed sequence-avalanches expressing high susceptibility to activity changes (Spearman, p = 0.0279; Fig. 4.f).

Finally, mapping the critical exponents along the branching parameter confirmed that as sequence-avalanches approach the critical point, their probability distribution slopes converge toward theoretical MFBP values. The duration exponent α converged to its expected theoretical value at criticality (Spearman, p < 0.001; Fig. 4.h). While the size exponent τ also correlated with proximity to criticality (Spearman, p = 0.03; Fig. 4.g), its absolute values exhibited a general offset lower than theoretically predicted. This systematic deviation is likely an artefact of temporal subsampling in calcium imaging, wherein large, distinct groups of neurons are aggregated as a single avalanche despite being separated by silent periods in reality (Capek et al., 2023; Neto et al., 2022; Srinivasan et al., 2024).

These findings establish that the brain might reside in a composition of heterogeneous states where overlapping neural substrates populate a phase transition line representing distinct sub- states at varying distances to criticality. Crucially, arousal-informed avalanches selectively operate near the critical point, characterized by high susceptibility and approximately scale-free dynamics.

## Methods

### Animals and surgery

All animal procedures and experiments were approved by the Institutional Animal Care and Use Committee at the University of California, Santa Barbara, and were carried out in accordance with the regulations and guidelines of the US Department of Health and Human Services. We used adult triple-transgenic mice expressing GCaMP6s in cortical excitatory neurons, generated by crossing TITL-GCaMP6s mice (Ai94, Allen Institute; Madisen et al., 2015), Emx1-Cre mice (Jackson Labs stock #005628), and ROSA:LNL:tTA mice (Jackson Labs stock #011008), as previously described (C. H. Yu et al., 2021). Mice were housed on a reverse 12-h light/12-h dark cycle with ad libitum access to food and water. A total of 10 recordings, comprising 5 spontaneous and 5 sensory-stimulus sessions, were obtained from 8 mice, with some mice contributing multiple recordings on different days.

At least one week before imaging, mice were surgically implanted with a chronic cranial window over the right visual cortex and a head-fixation implant. During surgery, mice were anesthetized with isoflurane (1.5-1.8%) and acepromazine (1.5-1.8 mg/kg body weight). Carprofen (5 mg/kg body weight) was administered before surgery and for 3 consecutive days after surgery. Body temperature was maintained during surgery, and the eyes were protected with ointment. The scalp overlying the right visual cortex was removed, and a custom head-fixing imaging chamber or headplate was mounted to the skull using cyanoacrylate-based glue and dental cement. A craniotomy was performed over the right visual cortex and sealed with a glass coverslip.

### Mesoscopic two-photon calcium imaging

Mesoscopic two-photon calcium imaging was performed at the University of California, Santa Barbara, using a custom Diesel2p mesoscope (C. H. Yu et al., 2021). Instrumentation and image acquisition were controlled with ScanImage software (Vidrio Technologies). Mice were awake and head-fixed during imaging. GCaMP6s-expressing neurons were imaged in layer 2/3 of the right visual cortex, at depths of approximately 150-200 µm from the pial surface. Fields of view were positioned to cover primary visual cortex (V1) and surrounding higher visual areas.

Imaging was performed over a 3 x 3 mm² field of view at cellular resolution. Movies were acquired at 2048 x 2048 pixels, corresponding to approximately 1.46 µm per pixel, with a frame rate of 7.5 Hz. Each imaging session lasted approximately 45 min. Laser power was typically 60-70 mW per 1.5 x 3 mm imaging path. Recordings were obtained during either spontaneous activity or visual-stimulation conditions, as described below.

### Brain region mapping

After recovery from cranial-window implantation, intrinsic signal optical imaging (ISOI) was performed to generate retinotopic maps and delineate visual cortical area boundaries, as previously described (Kalatsky & Stryker, 2003; Smith et al., 2017; Stirman et al., 2016; Y. Yu et al., 2022). Pial vasculature and intrinsic signal images were acquired through the cranial window using a CCD camera (Teledyne DALSA 1M30). A 4.7 x 4.7 mm² cortical region was imaged at 9.2 µm/pixel spatial resolution and 30 Hz frame rate.

Pial vasculature was illuminated and imaged through green filters (550 ± 50 nm and 560 ± 5 nm; Edmund Optics). Intrinsic signal images were collected by focusing 600 µm below the pial surface and illuminating/imaging through red filters (700 ± 38 nm, Chroma; 700 ± 5 nm, Edmund Optics). During ISOI, mice were lightly anesthetized with isoflurane (0.5%) and acepromazine (1.5-3 mg/kg), and body temperature was maintained at 37 °C using a custom electric heating pad (Stirman et al., 2016).

Visual stimuli for ISOI were presented on a flat monitor positioned 20 cm from the left eye. The monitor measured 60 x 34 cm² and covered approximately 110° x 75° of visual field. Responses to vertical and horizontal drifting bars were used to generate azimuth and elevation retinotopic maps. V1 and higher visual areas were identified from these maps, and area borders were manually drawn at azimuth and elevation meridians, following established criteria (Marshel et al., 2011; Smith et al., 2017). The vasculature map was then manually aligned to the two-photon maximum projection and used as a landmark to register retinotopically defined visual areas to the two-photon imaging field.

### Visual stimuli

Visual stimuli were presented to awake, head-fixed mice during passive viewing, with mice positioned either on a flat platform or on a floating treadmill allowing them to run freely. Stimuli were displayed on a 90 cm curved monitor positioned 14.5 cm from the mouse’s left eye and tilted 10° downward, covering approximately 150° of visual angle in azimuth and 70° in elevation. Visible emission from the screen was blocked from the imaging objective using a custom opaque plastic cone placed between the headplate and objective.

Sensory recordings consisted of randomized drifting grating patches. Grating patches were presented sequentially as 20°-wide square patches tiling the visual field. Stimulus parameters were randomized across presentations, including orientation, x-y position, spatial frequency, and temporal frequency, with parameter updates occurring at 2 Hz. Grating patches had a spatial frequency of either 0.04 or 0.1 cycles/degree and a temporal frequency of either 1 or 4 Hz, and were presented at 8 orientations: 0°, 45°, 90°, 135°, 180°, 225°, 270°, and 315°. The order of stimulus parameters was shuffled across successive presentations to enforce an even stimulus distribution. A projective correction was applied to preserve the intended stimulus geometry and position relative to the mouse eye.

Spontaneous recordings were obtained in the absence of patterned visual stimulation, either in total darkness or with the same screen set to a static middle-gray image.

### Pupil recording

Pupil size fluctuations were recorded during imaging using a custom-controlled infrared CCD camera (GigE, National Instruments) at 60 frames/s. The pupil was imaged through a small mirror positioned near and above the animal’s eye. No additional visible light stimulation was used for pupil imaging. Pupil diameters were extracted from the video recordings using DeepLabCut (Mathis et al., 2018; Nath et al., 2019).

### Calcium image data processing

Raw calcium imaging movies were first motion-corrected and aligned using the Suite2p registration module (Pachitariu et al., 2016). Regions of interest corresponding to individual neurons were then identified, and fluorescence traces were extracted using the EXTRACT algorithm (Dinç et al., 2021; Inan et al., 2017), which performs cell extraction while correcting for local background and neuropil contamination. Fluorescence traces were ΔF/F-normalized using EXTRACT.

Extracted traces were curated before downstream analysis. Cells were retained only if they had a signal-to-noise ratio (SNR) > 5, using the SNR estimate provided by EXTRACT, and cells located along the border of the field of view were excluded. After curation, recordings typically contained approximately 700-1000 cells/mm².

For avalanche detection and all subsequent analyses, we used neuronal activity traces that had been deconvolved with CASCADE, a deep-learning-based algorithm for estimating spike rates from calcium fluorescence signals (Rupprecht et al., 2021). The deconvolved spike-rate time series were used as the input to the avalanche-detection and seqNMF analyses described below.

### Neuronal avalanche detection

Neuronal avalanches were defined as temporally contiguous clusters of neuronal activity separated by silent periods (Beggs & Plenz, 2003). An avalanche corresponds to a contiguous sequence of non-empty imaging frames bounded before and after by empty frames. Avalanche size, S, was calculated as the total number of active neuronal events across all frames in the avalanche, corresponding to the sum of binarized population activity during the event. Avalanche duration, T, was calculated as the number of consecutive frames and converted to seconds using the imaging frame rate.

Avalanches were detected from deconvolved spike-rate time series using an optimizing- threshold approach (Bellay et al., 2015; Curic et al., 2021). For each recording, neuronal activity was represented as an N x T matrix, where N is the number of neurons and T is the number of frames. An optimized threshold, θ, was applied to the deconvolved activity of each neuron to obtain a binary activity matrix. The binary activity was then summed across neurons at each timebin to generate a thresholded population activity time series.

Because mesoscopic recordings from large neuronal populations do not contain silent frames in the unthresholded population signal, θ was selected separately for each recording to maximize the number of temporally separated avalanches. This procedure introduced empty time bins, where no neuron activity exceeded the threshold, allowing continuous population activity to be segmented into discrete avalanches.

### Power-law fitting and critical coefficients estimation

Avalanche size and duration distributions were analysed to estimate the scaling exponents τ and α, respectively. Size distributions were expressed as P(S) ∼ S^−τ^, where S is avalanche size, and duration distributions were expressed as P(T) ∼ T^−**α**^, where T is avalanche duration. Avalanche size was treated as a discrete variable, whereas avalanche duration was continuous in seconds.

For full-FOV avalanches, power-law slopes were evaluated over the scaling range displayed in Fig. 1: S > 10 for avalanche size and T > 1 s for avalanche duration. For sequence- avalanches, slopes were evaluated over the broader displayed scaling range used in Fig. 2 and Fig. 4: S > 10 for avalanche size and T > 2 s for avalanche duration. In each case, the exponent was estimated from the slope of the distribution in log-log space, with τ corresponding to the negative slope of the size distribution and α corresponding to the negative slope of the duration distribution.

For sequence-avalanches, exponents reported in the final figures were obtained from linear fits to the average probability-density curves over the displayed scaling ranges. Specifically, size exponents were fitted over S = 10-10,000, and duration exponents were fitted over T = 2- 20 s.

The duration cutoffs were chosen to be longer than the calcium indicator decay rate or the length of the sequence spatiotemporal motif.

### Sequence-avalanche decomposition

To identify overlapping spatiotemporal activity patterns that could not be separated by full-FOV avalanche detection alone, we decomposed the neuronal activity matrix using convolutional non-negative matrix factorization (seqNMF) (Mackevicius et al., 2019). seqNMF factorizes the activity matrix into a set of spatiotemporal motifs, W, and their corresponding temporal activation time series, H. Each W motif describes the neurons participating in a sequence and their relative timing within the motif, whereas H captures when and how strongly each sequence is expressed over the recording.

The number of candidate sequences, K, was empirically set to 20. The algorithm did prune any empty sequences. The maximum sequence duration, L, was set to 1.3 s, chosen to accommodate the temporal resolution of the mesoscope and the slow kinetics of the calcium indicator. This duration is also consistent with previous mesoscopic widefield imaging studies reporting sequence durations around 1 s (MacDowell and Buschman, 2020). The regularization parameter λ, which controls the trade-off between reconstruction error and orthogonality of the temporal and spatial factors, was iteratively adjusted until both reconstruction and orthogonality costs were minimized, following the approach described for seqNMF (MacDowell & Buschman, 2020; Mackevicius et al., 2019).

For each identified sequence, the corresponding W and H factors were convolved to reconstruct a sequence-specific activity matrix. Sequence-avalanches were then detected from these reconstructed activity matrices using the same optimized-threshold avalanche detection procedure described above. Sequence components were retained for downstream analysis if they explained more than 1% of the activity loading and contained more than 10 detected sequence-avalanches. This filtering yielded 62 valid sequence-avalanches across the 10 analysed datasets.

### Proximity to criticality

Proximity to criticality was quantified using the *d*_2_ distance metric described by Sooter et al. (2025). This method estimates how far an observed time series is from a critical dynamical regime without requiring direct knowledge of the underlying control parameter. The activity time series is first approximated by a vector autoregressive (VAR) model, which captures how current activity depends on recent past activity. The fitted autoregressive model is then compared with the set of autoregressive models that lie on the critical manifold, defined by Sooter et al. using a temporal renormalization-group framework. The *d*_2_ value is the minimum Kullback-Leibler divergence rate between the fitted model and the nearest model on the beta = 2 critical manifold. Thus, smaller *d*_2_ values indicate dynamics closer to criticality.

We used this metric because it provides a direct distance-to-criticality estimate from the activity time series itself, rather than relying only on avalanche size and duration power-law fits. This is useful for neural recordings, where the relevant control parameter is not directly accessible and may vary with behavioural or arousal state.

For full-FOV avalanches, *d*_2_ was computed from the thresholded avalanche time series for each recording. For sequence-avalanches, *d*_2_ was computed separately from each sequence- avalanche time series. Calculations were performed in MATLAB using the Sooter et al. codebase through the getDbetasTS function. The model order was set to 5, and the time step corresponded to the median sampling interval of each recording or sequence-avalanche time series. The returned d_beta value was used as *d*_2_.

### Branching parameters

The branching parameter, σ, was used to quantify how activity propagates from one time bin to the next. In an ideal branching process, σ corresponds to the average number of descendant active neurons generated by each active neuron. Values below 1 indicate subcritical dynamics, values close to 1 indicate near-critical dynamics, and values above 1 indicate supercritical dynamics (Haldeman & Beggs, 2005).

A direct branching-ratio estimate, computed from the ratio of sum of active neurons in consecutive time bins, can be strongly biased by spatial subsampling, because calcium imaging records only a subset of the full neuronal population. We therefore estimated σ using the multistep regression approach implemented in MR. Estimator (Spitzner et al., 2021). This method treats population activity as a branching-like autoregressive process and estimates how autocorrelation decays across multiple time lags. For such a process, the correlation coefficients follow *r_k_* = σ^k^, and the intrinsic timescale is τ = −Δt / ln(σ), where Δt is the sampling interval. Spatial subsampling mainly reduces the amplitude of the autocorrelation function, whereas the decay across lags is more robust; the MR estimator therefore provides a branching-parameter estimate that is less sensitive to recording only a fraction of the network.

For each full-FOV avalanche time series and each sequence-avalanche time series, autocorrelation coefficients were computed over multiple lags and fitted with an exponentially decaying function using the MR. Estimator toolbox. In our analyses, lags from 1 to 25 time steps were used, and the exponential-with-offset fit was used to obtain the MR estimate of σ.

### Information metrics

We use information-theoretic measures to first identify the arousal-related avalanches using mutual information and then characterise the dynamical properties of decomposed avalanches using information storage and Lempel-Ziv complexity.

The mutual information between the continuous sequence-avalanche timeseries and the corresponding pupil diameter was computed using the non-parametric KSG estimator in the Java Information Dynamics Toolkit (Lizier, 2014). Similarly, the information storage was also computed as the mutual information between the past k timesteps and the current timestep. We used *k* = 5 for this analysis.

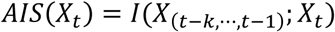

Here, *X_t_*is the timeseries of the sequence-avalanche and *X*_(_*_t_*_−*k*,⋯,*t*−1)_is the vector of past *k* timesteps. Lempel-Ziv complexity is computed using the LZ76 algorithm (Lempel and Ziv, 1976) applied on the binarized sequence-avalanche timeseries. We divided the timeseries into windows of length *L* = 1500 timesteps. The LZ76 algorithm returns the number of unique patterns within each window. The number of patterns was normalised to give an estimate of entropy rate ℎ as,

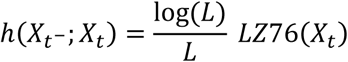

Dynamic range was computed from the range of avalanche sizes as 10 × log10(*S_max_*/*S_min_*), where *S_max_* and *S_min_* are the maximum and minimum avalanche sizes detected for a given sequence-avalanche. Shannon entropy was computed from the series of avalanche sizes and durations of each sequence-avalanche, yielding separate estimates of size entropy and duration entropy.

### Statistical physics metrics and phase diagram

Sequence-avalanches were positioned on a control-order phase diagram using the MR branching parameter, σ, as the control parameter. In this framework, σ < 1 indicates a subcritical regime in which activity tends to decay, σ ≈ 1 indicates proximity to the critical point, and σ > 1 indicates a supercritical regime in which activity tends to amplify. We then evaluated how macroscopic order parameters and fluctuation-based metrics varied as sequence- avalanches approached σ = 1.

Order parameters were computed from the sequence-avalanche population activity time series. Average activity, ρ, was defined as the mean fraction of active cells during avalanche activity. The normalized largest avalanche was calculated as the size of the largest detected avalanche divided by the number of neurons in the corresponding sequence. Synchrony was quantified as the mean sequence-avalanche population activity across the recording.

Susceptibility, χ, was computed as the variance-to-mean ratio of the sequence-avalanche activity time series: χ = = var(Aₜ)/mean(Aₜ), where Aₜ is the population activity at time bin t.

Conceptually, χ captures fluctuations in the order parameter; in second-order phase transitions, such fluctuations are expected to increase near the critical point.

Detrended fluctuation analysis (DFA) was used to quantify long-range temporal correlations in avalanche dynamics. DFA estimates a scaling exponent, α_DFA_, from the relationship between fluctuation amplitude and observation window size after local trends are removed. In general, α_DFA_ ≈ 0.5 indicates an uncorrelated random signal, α_DFA_> 0.5 indicates temporally correlated or persistent activity, α_DFA_≈ 1 indicates scale-free or 1/f-like long-range temporal correlations, and α_DFA_> 1 indicates very smooth or strongly persistent activity, which can occur when the signal contains slow rise-and-decay structure.

For full-FOV analyses, DFA was applied directly to the thresholded full-FOV avalanche time series using the nolds.dfa function from the Python nolds package for nonlinear-dynamics analysis.

For sequence-avalanche analyses, DFA was applied to an event-peak point process rather than to the raw continuous sequence-avalanche activity trace. This distinction was necessary because seqNMF represents each sequence as a continuous temporal activation profile. Even though the calcium traces were deconvolved before seqNMF, the seqNMF temporal factors and reconstructed sequence activity can still contain smooth activation envelopes, reflecting temporal jitter and variable expression strength across repeated sequence events. DFA applied directly to these smooth profiles did therefore yield inflated exponents, including values above 1, because the exponent is partly driven by the rise-and-decay shape of each event rather than by the temporal organization of avalanche occurrences.

To isolate avalanche timing, local maxima were detected in each sequence-avalanche activity time series using peak detection, with peaks required to exceed the mean activity of that sequence. Each sequence was then converted into a sparse binary point-process vector, with value 1 at detected avalanche peaks and 0 otherwise. DFA was applied to this binary event- peak time series, yielding an event-based DFA exponent, α_events_. This procedure removes the short-timescale autocorrelation introduced by smooth event shape while preserving the temporal spacing and clustering of avalanche occurrences.

## Discussion

This study demonstrates that neuronal avalanches in layer 2/3 of the awake mouse visual cortex operate near a critical point. By employing mesoscopic two-photon calcium imaging and convolutional non-negative matrix factorization (seqNMF) (Mackevicius et al., 2019), neural activity was decomposed into distinct, co-occurring avalanches. This dimensionality reduction approach revealed that the visual cortex is not uniformly positioned at a single distance from criticality. Instead, the system comprises multiple overlapping avalanches that populate the order-control phase diagram at various distances to the critical point, supporting the concept that low-dimensional spatiotemporal dynamics underlie widespread cortical activity (MacDowell & Buschman, 2020). Specifically, arousal-informed avalanches, which share mutual information with pupil diameter, operate close to the critical point and propagates widely from the posterior parietal and somatosensory cortices to the primary visual cortex. In contrast, uninformed avalanches tend to be localized and exhibit subcritical dynamics.

These findings address a significant gap in the literature by challenging the conventional assumption that neuronal avalanches occur as isolated events unique in time. By resolving concurrent sequence-avalanche dynamics, our results align with recent observations that scale-free, critical dynamics can be embedded within specific low-dimensional subspaces of high-dimensional awake brain activity (Fontenele et al., 2024). The observation of widespread, propagating pupil-linked avalanches is consistent with recent literature; arousal-elicited, brain-wide propagating waves have been observed on different recording modalities in mice and humans (Paz et al., 2026; Raut et al., 2021, 2025; Yang et al., 2026). Furthermore, the coupling of these avalanches to pupil dynamics supports existing evidence that pupil fluctuations and cortical desynchronization track fast switching of brain states to optimize sensory encoding (Reimer et al., 2014; Waschke et al., 2019).

Diverse cortical wave patterns have also been observed with voltage imaging (Liang et al., 2023). Functional ultrasound recordings revealed a common brain-wide hemodynamic wave during spontaneous and sensory-evoked arousal (Martinez de Paz et al., 2026, preprint), supporting consideration of arousal across stimulation contexts. Rajpal et al. (2026) reported longer spatial correlations and more extended synergistic interactions during visual stimulation.

Signatures of critical dynamics have now been reported in vivo across a wide range of brain states (Fontele et. al. 2025): during slow-wave and REM sleep (Scarpetta et. al., 2023, Priesemann et. al. 2013), in quiet wakefulness (Bellay et. al, 2015, Scott et. al., 2014), under sensory stimulation (Shew et. al. 2015), and during active task engagement (Yu et. al. 2017, Cambrinha et. al., 2025). Taken together, these studies point less to a disagreement about whether cortex is critical than to a system that remains in the neighbourhood of a critical point while its precise distance from that point shifts with state. Harder to establish has been the variable along which that distance is organised, particularly since these paradigms differ in species, preparation, recording modality, and choice of criticality estimator. By decomposing population activity into overlapping, sequence-specific avalanches, we find that arousal orders these sequences along the distances to criticality. Because arousal varies both within and across all of the paradigms above yet is seldom measured, it is a parsimonious candidate for the axis organising these otherwise heterogeneous reports. In future work, this hypothesis could be tested by replicating these conditions with concurrent pupillometry or another reliable index of autonomic arousal.

Pupil-linked activity in the locus coeruleus and in cortical noradrenergic and cholinergic axons provides candidate neuromodulatory pathways (Joshi et al., 2016; Costa and Rudebeck, 2016; Reimer et al., 2016). Pupil-associated VIP recruitment and SST suppression offer a potential cellular substrate (Reimer et al., 2014), although arousal effects differ across cell types (Kaufman et al., 2026).

Munn et al. (2026) linked pupil-inferred arousal to time-varying avalanche susceptibility in mouse spiking data and identified a directed-percolation transition in their model. Our sequence-resolved approach complements this work by distinguishing overlapping activity patterns at different distances from criticality, while retaining the temporal limitations of calcium imaging. These findings fit within the broader criticality framework reviewed by Hengen and Shew (2025).

Operating near the critical phase transition is theoretically predicted to maximize computational capabilities, including information capacity, fidelity of transmission, and dynamic range (Beggs, 2008; Shew et al., 2009; Shew & Plenz, 2013). The current data is consistent with aspects of this prediction, showing greater avalanche size dynamic range and lower Lempel-Ziv complexity within near-critical, arousal-informed avalanches (Mediano et al., 2023). Lower LZ complexity indicates that the analysed activity sequences contain more repeated patterns and are therefore more compressible. This is consistent with greater temporal regularity, although it does not by itself establish enhanced computational capacity. Furthermore, temporal entropy is maximized near the critical point, reflecting an expanded temporal state space and optimized long-term information storage necessary for handling tasks requiring both highly reactive processing and sustained working memory. This structural optimization likely serves a distinct mechanistic role, facilitating robust inter-area communication along the rostro-caudal axis and mediating transitions between exteroceptive and interoceptive modes (Yang et al., 2026; Raut et al., 2021), consistent with findings that cortical circuits are actively and homeostatically tuned to criticality in vivo to support cognitive tasks and adapt to structured sensory input (Habibollahi, Kagan, et al., 2023; Habibollahi, Sun, et al., 2023; Ma et al., 2019). From a clinical perspective, the segregation of these distinct dynamic regimes suggests that deviations from criticality could serve as highly specific biomarkers. Pathological shifts toward extreme sub-criticality or super-criticality within defined subnetworks may underlie network dysfunctions observed in disorders of consciousness or epileptic seizures (Gervais et al., 2023).

While mesoscopic calcium imaging provides an unprecedented spatial scale for avalanche detection, several methodological limitations constrain the interpretation of these scale-invariant properties. The seqNMF algorithm extracts repeated spatiotemporal templates of fixed maximum durations, which inherently limits the observation of true scale-invariance because it relies on convoluted temporal weight signals and cannot fully account for the unique spatial jitter or individual cellular participation present at each occurrence. Additionally, conventional avalanche detection relies heavily on the choice of temporal bin size and spatial subsampling. Similarly, recording from a limited number of neurons in a highly interconnected network can distort the observed scaling exponents. Finally, the slow kinetics of calcium indicators act as a temporal low-pass filter, aggregating underlying spiking activity, which may offset size exponent estimations from ideal mean-field branching process values. Our estimation of the branching parameter, for instance, generally remained below 1; while this indicates a subcritical regime on average, the non-stationary nature of awake cortical dynamics suggests that transient supercritical excursions likely occur and remain biologically relevant. Finally, Lempel-Ziv complexity should be interpreted cautiously. Although it provides a useful compression-based summary of temporal irregularity, recent work has shown that LZ- derived estimates of neural signal diversity can be strongly influenced by preprocessing choices, binarization, spectral structure, and the temporal scale at which the signal is analysed (Medel et al., 2023; Mediano et al., 2023). Future work should therefore complement the present LZ analysis with spectrally and temporally resolved signal-diversity estimates, as proposed by Mediano et al., to determine whether near-critical sequence-avalanches show enhanced complexity at specific timescales or frequency ranges.

Future investigations should link these mesoscopic statistical observations to the cellular and circuit mechanisms that generate and regulate near-critical avalanches, including the roles of specific cell types, synaptic interactions, and neuromodulatory pathways. The deployment of high-throughput high-speed voltage or calcium imaging (Bai et al., 2024; Chen et al., 2025; Manley et al., 2024) will be crucial to circumvent the temporal limitations of calcium indicators and accurately resolve the fine-scale temporal structures of overlapping avalanches. Moreover, while this study establishes a correlation between arousal and near-critical network dynamics, causal perturbational experiments, such as optogenetic (Ribeiro et al., 2026) or pharmacological interventions (arousal manipulation) targeting neuromodulation (Stewart and Plenz, 2006), are required to determine whether targeted manipulations can actively tune specific subnetworks into or out of criticality and directly alter behaviour. It would similarly be interesting to manipulate brain criticality through imposed behaviours, building on observed links between scale-free behavioural and cortical dynamics (Jones et al., 2023). Understanding how co-occurring avalanches support specific cognitive processes, and how scale-free cortical dynamics relate to behaviour, remains central to establishing the functional role of criticality in the brain. Our findings motivate combining pupillometry with non-invasive brain recordings, such as EEG, fMRI, or functional near-infrared spectroscopy (fNIRS), to investigate the relationship between arousal, criticality, and cognitive performance. Together, these measurements could help determine whether criticality provides a useful biomarker of cognitive function and its disruption in psychiatric and neurodegenerative disorders (O’Byrne & Jerbi, 2022; Zimmern, 2020).

## Acknowledgments

We thank Lucilla de Arcangelis for her participation in the Statistical Physics of Cognition project and for helpful discussions during the early stages of this work.

## Author contributions

C.S. conceived the study, processed and analysed the data, prepared the figures, and wrote the original manuscript. H.R. computed information-theoretic metrics and contributed to scientific discussions and interpretation. J.C. performed mouse surgeries and calcium-imaging experiments and generated the experimental data. M.G.K. performed calcium-signal deconvolution. M.S., H.J.J., and M.B. contributed to scientific discussions and interpretation, with H.J.J. providing expertise in avalanches and criticality and M.B. in network science. S.L.S. supervised experimental data collection. S.R.S. led the grant and overall project, provided overall supervision, and contributed to scientific discussions and interpretation. H.R., J.C., M.S., H.J.J., M.B., S.L.S., and S.R.S. reviewed the manuscript.

