## Supplemental Figures for "Arousal-informed neuronal avalanches operate near criticality"

### Supplementary figures

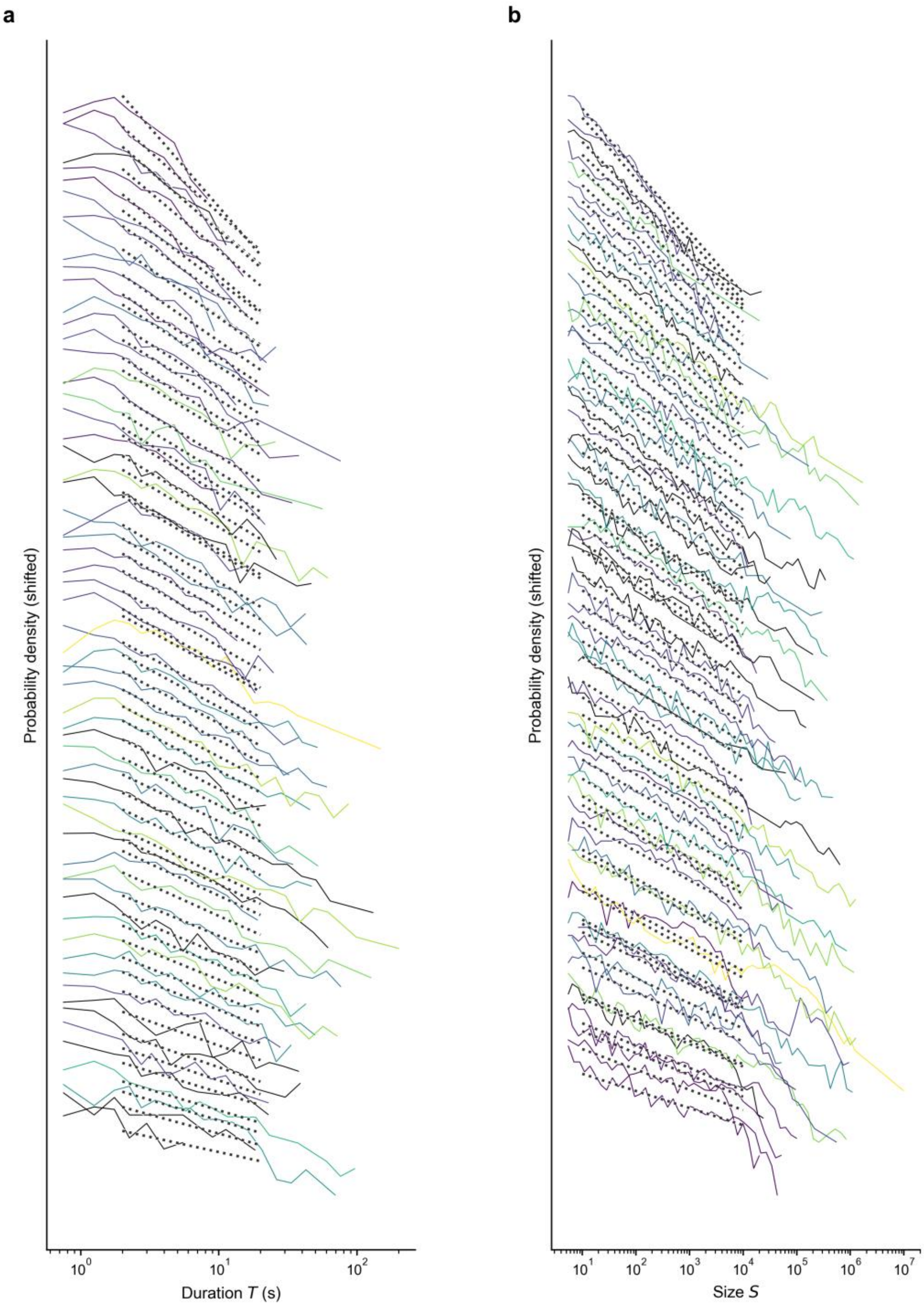


**Figure S1. Individual sequence-avalanche duration and size distributions.**

**(a, b)** Probability density distributions for avalanche duration (T, panel a) and size (S, panel b) for individual sequence-avalanches. Distributions are vertically shifted for clarity. Colored solid lines indicate empirical distributions, color-coded by mutual information (MI) with pupil diameter, from low MI (purple) to high MI (yellow), as in Figure 2. Black solid lines indicate sequences without available pupil recordings. Black dotted lines indicate the individual power-law fits used to estimate the duration exponent α and size exponent τ.

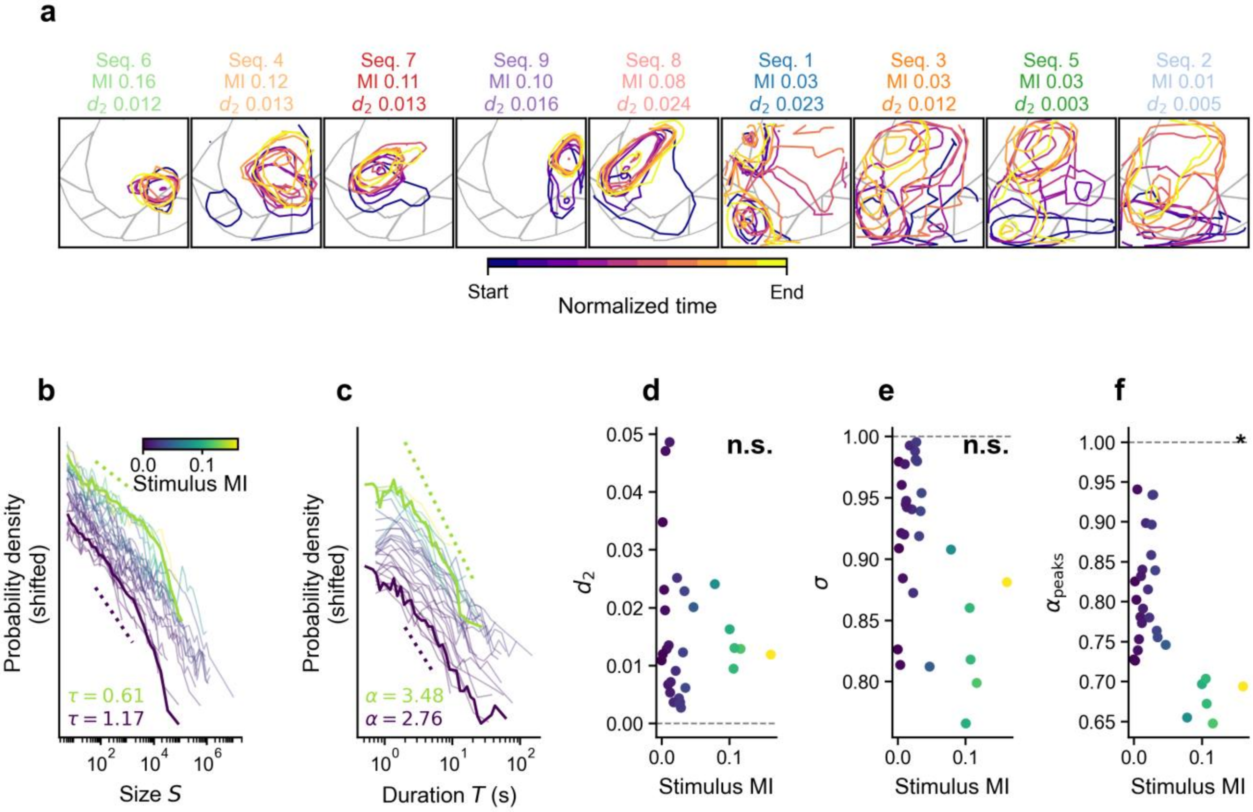


#### Figure S2. Sequence-avalanche dynamics as a function of stimulus mutual information.

**(a)** Spatiotemporal footprints of nine sequence-avalanches, ordered by their mutual information (MI) with the visual stimulus. Sequence labels show stimulus MI and proximity to criticality ($d_2$). Colors of the contours indicate normalized time from start (purple) to end (yellow) of the sequence motif. **(b, c)** Probability density distributions for avalanche size (S, panel b) and duration (T, panel c) for individual sequences, color-coded by stimulus MI. Distributions are vertically shifted for clarity. Thick solid lines show the average distributions for sequences below the 15th percentile and above the 85th percentile of stimulus MI; dotted lines indicate the corresponding power-law fits, with size ($\tau$) and duration ($\alpha$) exponents displayed. **(d-f)** Scatter plots show **(d)** proximity to criticality ($d_2$), **(e)** branching parameter ($\sigma$), and **(f)** event-peak DFA exponent ($\alpha_{peaks}$) plotted against stimulus MI for each sequence. Dashed horizontal lines mark $d_2 = 0$, $\sigma = 1$, and $\alpha_{peaks} = 1$, respectively. Only sequences with available stimulus MI are included (n = 29 sequences from four stimulated recordings); stimulus timing and MI were unavailable for mouse3. Statistical significance was assessed using Spearman's rank correlation test: *$p < 0.05$; n.s., not significant. Stimulus MI was negatively associated with event-peak DFA ($r_s = -0.383$, $p = 0.040$), but not significantly associated with $d_2$ ($r_s = -0.148$, $p = 0.444$) or $\sigma$ ($r_s = -0.273$, $p = 0.152$).
